# Benchmarking datasets for machine learning in protein function prediction

**DOI:** 10.64898/2025.12.17.694800

**Authors:** Yaan J. Jang, Qi-Qi Qin, Jin-Lei Wang, Benoît Kornmann

## Abstract

Remarkable progress has been achieved by machine learning, particularly in accurate prediction of protein tertiary structures. Despite these advances, accurately annotating protein functions through machine learning approaches remains challenging, primarily due to the limited availability of large-scale benchmarking data. In this study, we addressed this gap by systematically screening proteins from the UniProt database for functional annotations, resulting in the creation of a benchmarking dataset that includes protein sequences and their corresponding annotations. The Protein Annotation Dataset (PAD) is a resource available to train a wide range of machine learning models for assignment of function annotations to previously unlabeled proteins. We curated a comprehensive dataset comprising four categories of functional annotations using enzyme commission (EC) numbers and gene ontology (GO) terms. The dataset was subsequently partitioned into training, validation, and test subsets. Furthermore, we incorporated an independent set from 12 diverse species, enabling the development and evaluation of innovative machine learning models.

## Background & Summary

Understanding protein function is crucial for deciphering the complexities of biological processes and their implications for health and disease. Genomes record billions of evolution experiments, yielding myriads of variants across a multitude of protein sequences. These variants, numbering in the millions, dictate the suitability of proteins for specific functional roles in biological activities. With the advent of efficient and cost-effective sequencing technologies, machine learning (ML), particularly deep learning models, has emerged as a powerful tool for extracting meaningful insight from these sequences.

However, while significant progress has been made in utilizing machine learning to predict protein tertiary structures^1–4^, predicting protein function remains challenging, especially for protein families lacking comprehensive functional annotations. Indeed, state-of-the-art approaches assign functional annotations to proteins by leveraging either structural data or homologous information^5^, ^6^. They often rely on assuming that proteins with significant sequence similarity share similar functions^7–9^. While effective, these approaches have limitations, particularly with distantly related or highly divergent proteins, and are not suitable for entirely novel proteins lacking close homologs.

As a promising alternative, ML-based methods have been developed to predict protein function annotations^5, 6^, ^10^, including gene ontology (GO) terms^11^, which provide standardized descriptors of protein functions and localization, as well as enzyme commission numbers (ECN) annotating enzyme-catalyzed reactions^12^. These methods can capture intricate knowledge within protein sequences and structures, enabling accurate function prediction even in the absence of closely related homologs. Despite significant progress, ML-based methods for protein function prediction require diverse training datasets encompassing vast volumes of data from various sources^6^, ^13^. With the exponential growth of sequence data, there is a need for benchmark datasets to enhance the efficiency and accuracy of these methods. Many proteins remain poorly characterized, and even well-studied proteins may have incomplete functional annotations^10^, particularly challenging when generalizing from known proteins to novel ones, especially from non-model organisms, as their functions may differ substantially from those of well-studied model organisms.

High-quality and diverse training data significantly enhance the accuracy of deep learning models for protein function prediction^6, 14^, ^15^. To advance ML-based methods, we have compiled extensive protein data from the UniProt database^16^, establishing a robust foundation for functional annotations. The Protein Annotation Dataset (PAD) provides a comprehensive resource for training and validating models. This endeavour represents a significant advancement for understanding protein function through machine learning, with applications ranging from drug discovery to personalized medicine.

## Methods

In this section, we outline the process of collecting data, including our criteria for inclusion and the subsequent filtering steps. Moreover, we present detailed statistics to enhance the dataset’s usability. We also highlight the potential of the PAD dataset for the development of ML-based solutions, discussing its applicability. Finally, we acknowledge and address any potential limitations inherent in the dataset to provide a comprehensive overview.

### Data collection

In this study, we curated benchmark datasets tailored for ML-based protein function annotation, encompassing both ECNs and GO terms that compose of biological processes (BP), cellular components (CC), and molecular functions (MF). Our datasets were constructed using the UniRef50 database within UniProt^16^ (released on June 28, 2023). This database contains 60,952,894 protein sequences, clustered using a 50% sequence identity threshold. Of these sequences, 13,290,382 are identified as enzymes based on their nomenclature, characterized by names ending with ‘ase’^17^. Our data collection process began by retrieving the UniProt IDs (UIDs) for all proteins in the UniRef50 database. We then developed an in-house script to systematically screen each protein and query whether it had been functionally annotated in the UniProt database^16^. Utilizing these UIDs, we accessed the UniProt web-server to retrieve protein entries in the JavaScript Object Notation (JSON) format. Leveraging these JSON files, we extracted function annotations for approximately 60 million proteins spanning BP, CC, and MF, along with approximately 13 million enzymes annotated with ECN.

Our thorough examination of the raw data has uncovered significant insights: a vast majority of proteins remain unannotated, lacking crucial functional information. Specifically, our analysis reveals that over 75% of proteins in the raw dataset lack any GO annotations, severely limiting our understanding of their roles and interactions within biological systems. Similarly, approximately 80% of the staggering 13 million enzymes analyzed remain unlabeled with ECNs, indicative of a significant gap in our knowledge of enzymatic functions (Fig. 1). These findings underscore the importance of continued efforts in protein function annotation, particularly in bridging the gap between annotated and unannotated proteins. While the availability of meticulously curated annotations is significant, there exists an urgent imperative to broaden coverage to encompass all proteins, especially enzymes, to unlock the full potential of functional genomics. Furthermore, our analysis highlights the diversity and complexity of protein functions, as evidenced by the multitude of unique labels identified across various functional categories.

**Figure 1.**
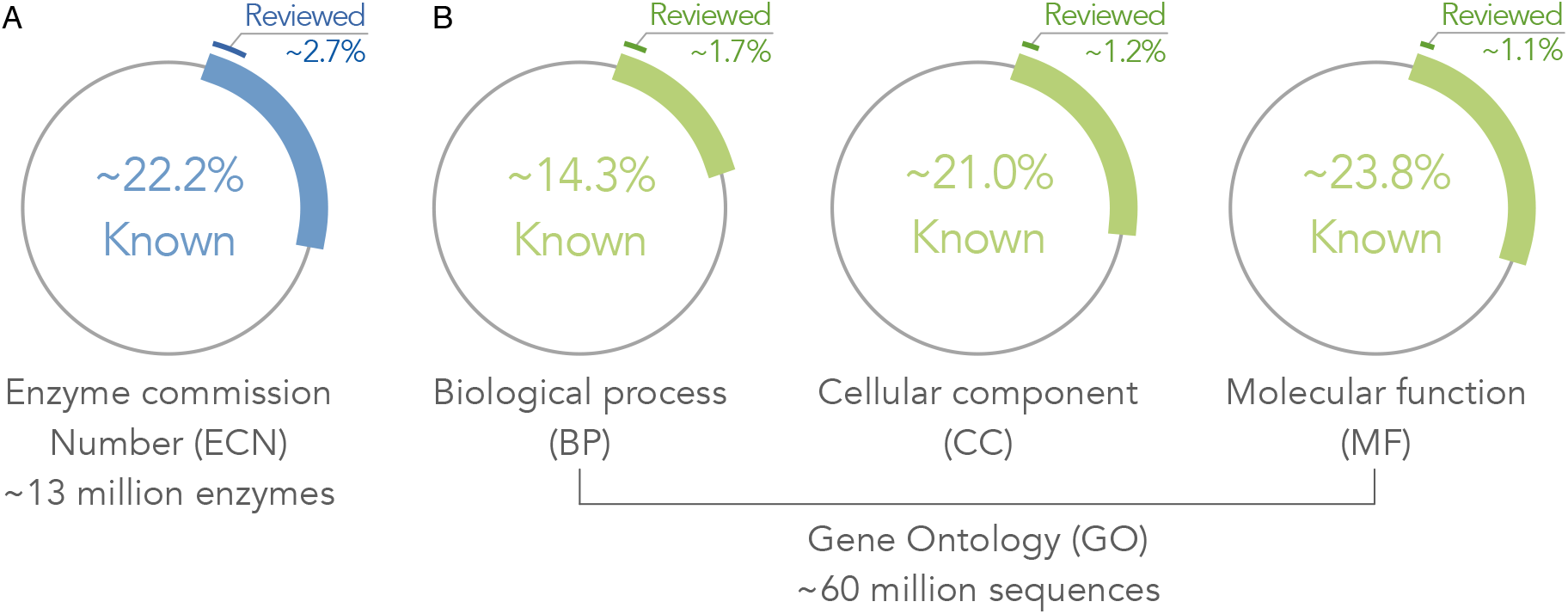
Statistics of proteins in the UniRef50 database. (A) Enzymes of known ECN. About 22% of the approximately 13 million enzymes are annotated with ECN, of which less than 3% of the annotated enzymes are expertly reviewed. (B) Proteins of known GO terms. The expertly reviewed GO terms are all less than 2% of the proteins with annotated functions.

The proteins that do have known annotations are classified into two distinct categories: reviewed proteins, meticulously curated by dedicated biocurators, and unreviewed proteins, which have not undergone human curation. Our examination of the dataset has identified approximately 3 million enzymes annotated with ECN, illustrating a partial but significant coverage in this domain (Fig. 1)A). Additionally, we observed annotations for approximately 9 million proteins in terms of BP, 13 million proteins in terms of CC, and 15 million proteins in terms of MF (Fig. 1)B, and Table 1). In total, our obtained dataset encompasses a diverse array of labels, providing valuable insights into protein function annotation. Specifically, we have identified 6,053 unique labels for ECN, 18,234 for BP, 2,842 for CC, and 8,334 for MF. Remarkably, these labels are further stratified based on the reviewed status of the proteins, offering a nuanced understanding of the annotation landscape. Notably, 5,128 labels for ECN, 17,772 for BP, 2,772 for CC, and 8,029 for MF are attributed to reviewed proteins, indicating a substantial presence of expertly curated annotations within our dataset. However, despite the prevalence of reviewed annotations, our analysis reveals a concerning disparity: less than 3% of annotated proteins undergo expert review. This leaves the majority of annotated proteins susceptible to inaccuracies or incompleteness, particularly within the unreviewed category. This disparity underscores the urgent need for comprehensive annotation efforts, combining automated methods with rigorous manual curation to enhance the accuracy and completeness of protein annotations.

**Table 1.** Proteins of annotated functions obtained from the UniRef50 database.

| Category | #proteins |  |  |  | #unique labels |
| --- | --- | --- | --- | --- | --- |
|  | Total | Annotated | Reviewed | Unreviewed |  |
| ECN | 13,290,382 | 2,952,970 | 79,371 | 2,873,599 | 6,053 ( 5,128*) |
| BP | 60,952,894 | 8,741,373 | 149,821 | 8,591,552 | 18,234 (17,772*) |
| CC | 60,952,894 | 12,810,123 | 148,917 | 12,661,206 | 2,842 ( 2,772*) |
| MF | 60,952,894 | 14,528,998 | 153,957 | 14,375,041 | 8,334 ( 8,029*) |

\*Total number of the unique labels from the reviewed proteins.

### Data filtering

In ML-based models, particularly those utilizing deep neural networks, the number of parameters often aligns with the maximum number of residues present. This correlation leads us to establish a threshold for the number of residues in proteins, aiming to strike a balance between the number of trainable parameters and the coverage of proteins. To thoroughly evaluate this threshold, we categorized proteins into the four distinct groups based on their reviewed and unreviewed statuses. This categorization facilitated an examination of how protein length fluctuates between curated and non-curated records (Fig. 2).

**Figure 2.**
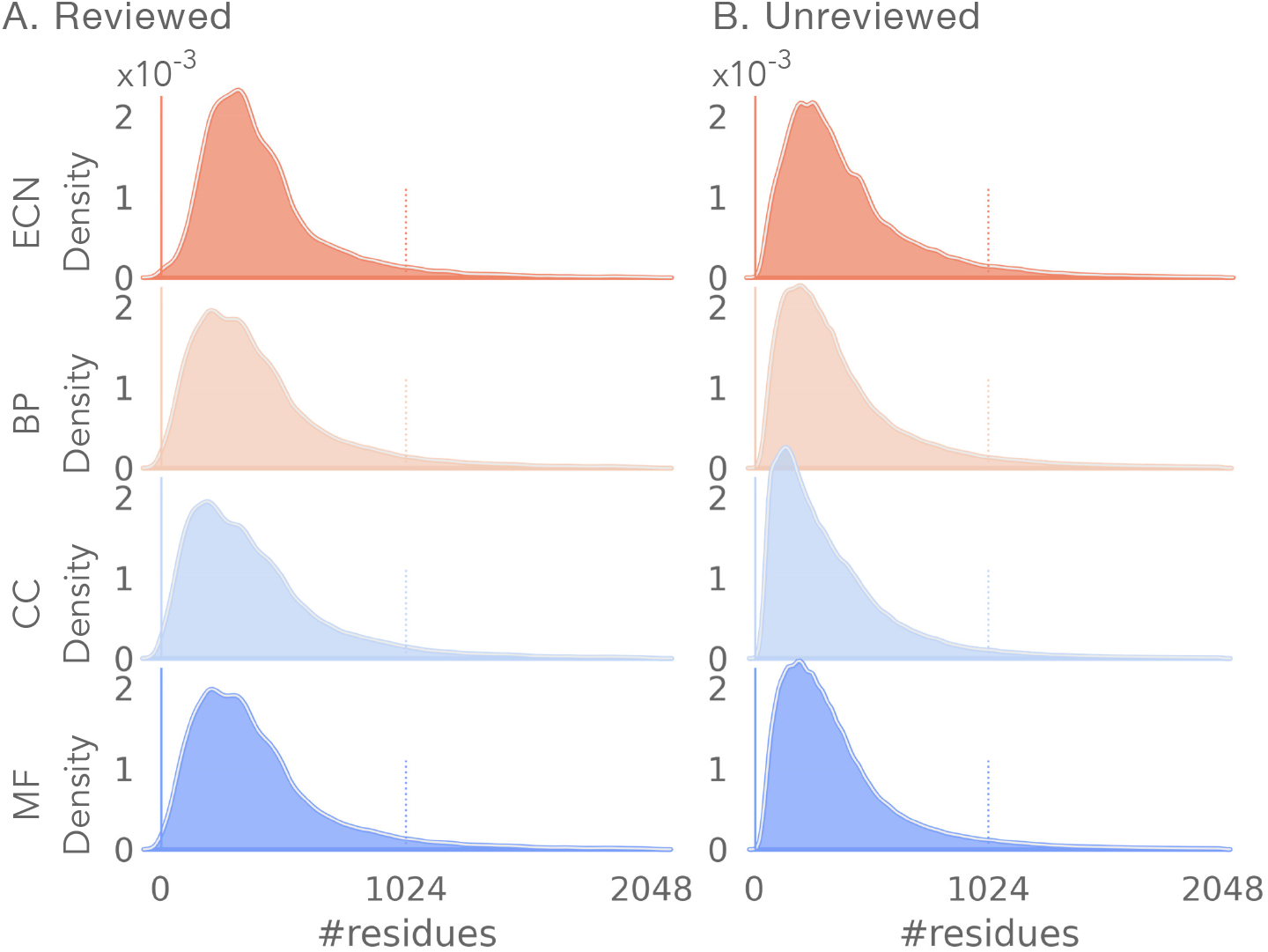
The distributions for the number of residues in proteins with (A) reviewed and (B) unreviewed function annotations.

Our analysis focused on proteins containing up to 2,048 residues, excluding those with longer sequences due to their relatively minor representation in the dataset. The majority of proteins in both reviewed and unreviewed categories exhibited a length of fewer than 1,024 residues, indicating a shared pattern in sequence length distribution across both groups. Across all functional categories (ECN, BP, CC, and MF), the distributions of protein lengths exhibited similar shapes for both reviewed and unreviewed proteins. Nevertheless, the CC category exhibited marginally more pronounced disparities in length distribution compared to the other three categories.

This scrutiny underscores the criticality of selecting the maximum number of residues when devising machine learning models for protein function prediction. While subtle variations exist in length distributions between reviewed and unreviewed proteins, the overarching similarities imply that model architectures should possess robustness to accommodate diverse sequence lengths across various functional categories. Moreover, our findings highlight the necessity for focused attention on protein length, particularly within the CC category where discrepancies in length distribution are more apparent. The development of ML algorithms capable of effectively capturing these nuances in length distribution, while preserving model efficiency and accuracy, is indispensable for advancing protein function prediction methodologies. In light of these findings, we advocate for a careful consideration of protein length when designing ML-based models for protein function prediction. While there are subtle differences in length distributions between reviewed and unreviewed proteins, the overarching similarities suggest that model architectures should be robust enough to accommodate varying sequence lengths across different functional categories. Additionally, special attention should be paid to functional categories where differences in length distribution are more apparent, such as the cellular component category.

### Data generation

The size of proteins plays a pivotal role in the realm of ML-based methods. Proteins of more than 1,024 residues would necessitate more intensive computational resources for both training and prediction tasks, especially in expansive applications. While proteins with fewer residues may not offer sufficient information to effectively train machine learning models, potentially limiting their predictive capabilities. To address this, we implemented three filtering rules to refine our dataset: (1) exclusion of proteins by length, protein sequences with less than 32 or more than 1,024 amino acids were excluded from the dataset to focus on proteins of intermediate length, which are more likely to offer informative features for model training, yet not overly burdensome in terms of computational resources; (2) exclusion of unreviewed labels of function annotations, annotations associated with unreviewed proteins were systematically excluded to ensure the reliability and quality of the annotations used for model training and evaluation; and (3) annotation filtering, annotations assigned to fewer than 16 proteins were removed from the dataset, aiming to retain annotations with sufficient representation to support robust model training and prediction. By implementing these filtering rules, we obtained a refined dataset comprising proteins and their annotations, as detailed in Table 2. In the filtered dataset, the number of unique annotations (labels) varies depending on the specific filtering criteria applied. This variability directly influences the total number of parameters in the ML-based models, with fewer unique annotations leading to reduced models’ predictive ability. To explore the impact of varying the number of proteins per unique annotations over the number of the annotations, we conducted calculations with different thresholds (N=1, 16, and 32) to filter proteins.

**Table 2.** Proteins of *L* ∈ [32, 1024] and reviewed labels are filtered from the UniRef50 database.

| Category | #proteins (reviewed) |  |  | #proteins (unreviewed) |  |  | #unique labels |  |  |
| --- | --- | --- | --- | --- | --- | --- | --- | --- | --- |
| | $N^{\dagger} \geq 1$ | $N \geq 16$ | $N \geq 32$ | $N \geq 1$ | $N \geq 16$ | $N \geq 32$ | $N \geq 1$ | $N \geq 16$ | $N \geq 32$ |
| ECN | 72,465 | 68,878 | 67,896 | 2,722,210 | 2,716,510 | 2,709,244 | 5,033 (6,010*) | 3,131 | 2,802 |
| BP | 139,662 | 129,042 | 123,358 | 8,094,836 | 8,081,320 | 8,066,503 | 17,221 (17,221*) | 6,685 | 4,736 |
| CC | 138,392 | 134,955 | 133,538 | 12,112,921 | 12,109,695 | 12,106,581 | 2,682 (2,784*) | 1,387 | 1,155 |
| MF | 144,445 | 138,968 | 136,579 | 13,705,138 | 13,697,009 | 13,686,475 | 7,830 (8,231*) | 4,529 | 3,876 |

<sup>†</sup>Number of proteins per label.
\*Total number of the unique labels, including reviewed and unreviewed labels.

**Table 3.**
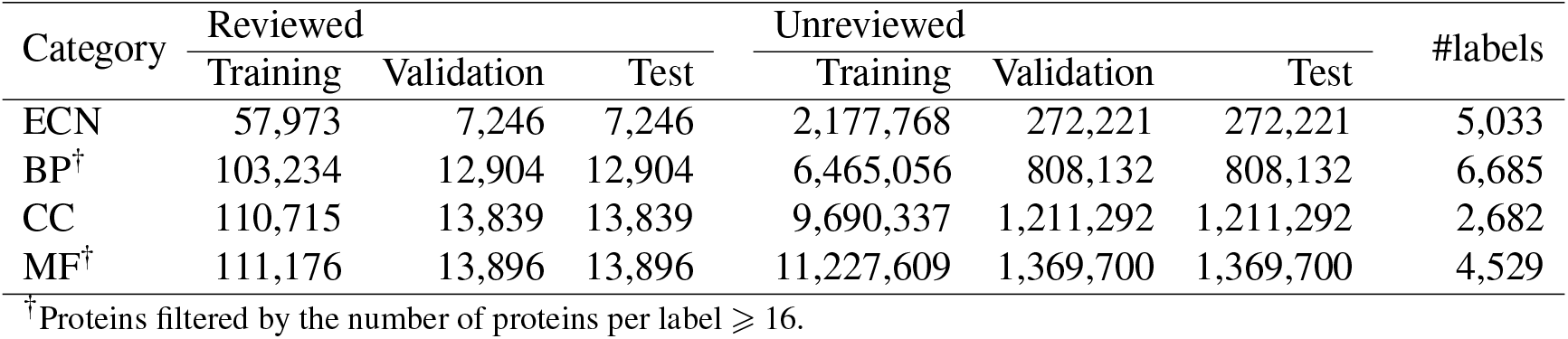
Benchmark datasets of protein function annotations.

This strategy to filtering ensures that our dataset strikes a balance between computational efficiency and the richness of information available for model training and prediction. By systematically refining the dataset based on the rules, we aim to help optimize model performance while maintaining the integrity and relevance of the underlying protein annotations. These filtered datasets would be valuable resources for developing and fine-tuning ML models in protein function prediction.

### Benchmark datasets for machine learning

To enhance the predictive performance of ML-based models for protein function prediction, we embarked on constructing comprehensive benchmark datasets tailored specifically for training and evaluating these models. Our approach involved meticulously curating two distinct benchmark datasets: one comprising reviewed proteins, meticulously curated by biocurators, and the other encompassing unreviewed proteins. Achieving a balance between model performance and complexity necessitated the implementation of a meticulous label filtering process. To ensure that our datasets contained labels with a sufficient number of data points for robust analysis, we applied a stringent criterion: labels associated with functional categories containing fewer than 16 proteins in the categories of BP and MF were excluded from the dataset. This filtering was crucial in maintaining the integrity and reliability of the labels used for model training and evaluation. Following the label filtering procedure, our datasets retained a total of 5,033 labels for ECN, 6,706 labels for BP, 2,682 labels for CC, and 4,560 labels for MF. The meticulous curation process ensured consistency in labels across both the reviewed and unreviewed protein datasets, thereby facilitating direct comparisons between machine learning models trained on these datasets. This consistency was pivotal in enabling a comprehensive assessment of model performance and generalization across diverse protein datasets.

Subsequently, we partitioned each dataset into three distinct subsets: training, validation, and test sets, in a ratio of 80%:10%:10% respectively. This strategic division of the datasets was designed to provide abundant data for model training while ensuring sufficient data for rigorous model validation and test. By adopting this partitioning strategy, we aimed to strike a balance between model complexity and generalization, thereby optimizing model performance across various evaluation metrics. The meticulous construction of these benchmark datasets represents a significant milestone in the field of protein bioinformatics and machine learning. These datasets serve as a reliable foundation for the development and evaluation of ML-based models and tools aimed at elucidating the intricate functionalities of proteins in diverse biological processes.

### Independent hold-out test datasets of species-specific proteins

Indeed, when ML models are presented with proteins from diverse species during evaluation, they are effectively put to the test on their capacity to predict function annotations for proteins that may exhibit unique structural and functional characteristics not previously encountered during training. This rigorous evaluation ensures that ML models demonstrate robustness and versatility in their predictive capabilities, thereby instilling confidence in their applicability across a wide range of biological contexts and taxonomic groups. Furthermore, the independent test dataset is a valuable resource for the broader scientific community, facilitating benchmarking and comparison of different ML models and methodologies in the field of protein function prediction. By providing a standardized evaluation framework grounded in real-world biological data from diverse species, the independent test dataset contributes to advancing the state-of-the-art in machine learning-based protein function prediction. Our selection of these species was purposeful and strategic, aiming to encompass a broad spectrum of organisms representing various taxonomic groups. This deliberate inclusion of both well-studied model organisms and less-explored species ensures that the independent test dataset presents a formidable challenge for ML methods, prompting them to make accurate predictions for proteins with potentially distinct functional characteristics across different taxa.

In the present work, we collected an independent test datasets from twelve species (Table 4). In our endeavor, the acquisition of an independent test dataset from twelve distinct species has been a pivotal undertaking, crucial for advancing the development and evaluation of innovative ML methods in the realm of protein function prediction. The process of constructing the independent test dataset involved meticulous curation of data from twelve diverse species (released by 2023/7/19) sourced from the UniProt database. Using the above protocol utilized for the construction of the training/validation/test benchmark datasets, we curated the independent test dataset with stringent adherence to established guidelines and criteria. This approach ensures the reliability and integrity of the dataset, setting the stage for rigorous evaluation of ML models’ generalization capabilities. The independent test dataset is a critical benchmark for assessing the performance and generalization capabilities of machine learning models trained on benchmark datasets comprising reviewed and unreviewed proteins. By evaluating models on proteins from diverse species not encountered during the training phase, the independent test dataset provides a comprehensive assessment of models’ ability to extrapolate functional annotations to novel proteins.

**Table 4.** Species-specific proteins of annotated functions (released by 2023/7/19).

| Organism | Unannotated enzymes<br>(w/ -ase suffix) | Annotated ($L \in [32, 1024]$) | | | |
| --- | --- | --- | --- | --- | --- |
|  |  | ECN | BP | CC | MF |
| Homo sapiens | 193,969-50,234 | 50,234 | 1,074,348 | 817,827 | 1,061,039 |
| Rattus norvegicus | 17,732-7,624 | 7,624 | 40,425 | 44,203 | 41,430 |
| Mus musculus | 21,557-10,916 | 10,916 | 52,579 | 66,671 | 68,311 |
| Arabidopsis thaliana | 67,659-27,907 | 27,907 | 99,235 | 102,624 | 130,336 |
| Danio rerio | 11,425-6,567 | 6,567 | 28,882 | 31,573 | 31,009 |
| Caenorhabditis elegans | 140,179-51,127 | 51,127 | 121,590 | 127,751 | 187,292 |
| Saccharomyces cerevisiae S288C | 2,072-1,690 | 1,690 | 5,312 | 6,047 | 4,634 |
| Drosophila melanogaster | 63,049-38,109 | 38,109 | 124,748 | 140,881 | 152,713 |
| Dictyostelium discoideum | 5,849-2,477 | 2,477 | 11,034 | 13,128 | 11,688 |
| Schizosaccharomyces pombe | 2,055-1,569 | 1,569 | 5,190 | 5,871 | 4,672 |
| Escherichia coli | 497,534-272,317 | 272,317 | 479,843 | 404,471 | 659,393 |
| SARS-CoV-2 | 399-211 | 211 | 8,453 | 9,303 | 5,791 |

Table 4 presents a summary of species-specific proteins along with their annotated functions, as of the release date of July 19, 2023. Meanwhile, Table 5 provides a summary of sequence clusters at 50% sequence identity for the species-specific proteins, generated using the MMseqs2 tool^18^. The species-specific proteins originate from a diverse array of organisms, spanning humans, bacteria, and viruses. Each characterized by varying counts of annotated proteins falling within a specific length range of 32 to 1024 amino acids. We categorized the proteins of known either ECNs or GO terms across the twelve species. As shown, Homo sapiens is a rich and well-characterized proteome, with a high count of annotated proteins across all functional categories. This reflects the extensive research focus on human biology and the importance of understanding human physiology and disease mechanisms. Model organisms like Mus musculus, Arabidopsis thaliana, and Caenorhabditis elegans also demonstrate substantial counts of annotated proteins, underscoring their utility in biological research as representative models for studying various biological processes. Microorganisms such as Escherichia coli and Saccharomyces cerevisiae S288C possess smaller proteomes compared to multicellular organisms but are characterized by a significant number of annotated proteins. This highlights the importance of studying microbial metabolism and cellular processes, with potential applications in biotechnology and industrial processes. Moreover, SARS-CoV-2, the virus responsible for the COVID-19 pandemic, is included. Despite its smaller proteome, SARS-CoV-2 possesses a notable number of annotated proteins, reflecting ongoing research efforts to understand viral pathogenesis and develop therapeutic interventions.

**Table 5.** Species-specific proteins of annotated functions clustered with 50% sequence identity. Organism Unannotated enzymes Annotated (*L* ∈ [32, 1024])

| Organism | Unannotated enzymes<br>(w/ -ase suffix) | Annotated ($L \in [32, 1024]$) | | | |
| --- | --- | --- | --- | --- | --- |
|  |  | ECN | BP | CC | MF |
| Homo sapiens | 193,969-50,234 | 6,570 | 40,719 | 40,144 | 43,981 |
| Rattus norvegicus | 17,732-7,624 | 3,033 | 16,894 | 18,085 | 16,150 |
| Mus musculus | 21,557-10,916 | 4,731 | 23,272 | 25,896 | 24,438 |
| Arabidopsis thaliana | 67,659-27,907 | 6,916 | 25,718 | 29,020 | 33,146 |
| Danio rerio | 11,425-6,567 | 2,948 | 13,103 | 14,266 | 12,970 |
| Caenorhabditis elegans | 140,179-51,127 | 10,683 | 34,500 | 4,4493 | 54,800 |
| Saccharomyces cerevisiae S288C | 2,072-1,690 | 1,483 | 4,741 | 5,426 | 4,054 |
| Drosophila melanogaster | 63,049-38,109 | 7,916 | 26,981 | 31,330 | 33,629 |
| Dictyostelium discoideum | 5,849-2,477 | 1,619 | 6,953 | 8,648 | 7,258 |
| Schizosaccharomyces pombe | 2,055-1,569 | 1,451 | 4,521 | 5,053 | 3,993 |
| Escherichia coli | 497,534-272,317 | 30,098 | 41,344 | 39,445 | 68,274 |
| SARS-CoV-2 | 399-211 | 103 | 480 | 521 | 433 |

In summary, the acquisition and curation of the independent test dataset (Table 4) from twelve diverse species offers valuable dataset for evaluating machine learning models’ generalization capabilities. Moreover, these models built on the dataset can advance our understanding of protein function and biological processes. Moving forward, the utilization of this independent test dataset will continue to drive innovation and foster advancements in the field of protein bioinformatics and machine learning.

### Data Records

All proteins and their function annotations in PAD were collected from both unreviewed and expertly reviewed records, and a website ^†^ provides tutorials for users to download the proteins and their function annotations, where the raw and pre-processed data are hosted.

The raw dataset contains ∼3 million enzymes of ECNs and ∼36 million proteins of GO terms. This dataset can be alerted for training and evaluating specific developments of ML-based models. The pre-processed datasets, including training, validation, test, and independent test datasets, are created for both pre-trained ML models and newly developed models.

Protein records and their function annotations are organized into separate folders, each containing a list of proteins categorized by function. For instance, the file “list_proteins_of_ECN_annotations.csv” contains proteins with known ECN annotations. The UIDs in this file facilitate matching protein sequences with their function annotations. Each protein record follows a format outlined in Table 6, featuring fields including “ID” of UniProt Identifier, “Name” of proteins in UniProt, “Status” of unreviewed/expertly reviewed, “#Residues” within each protein, and “Annotation” of each protein.

**Table 6.** Example data records of proteins with ECNs.

| ID | Name | Status | #Residues | Sequence | Annotation |
| --- | --- | --- | --- | --- | --- |
| Q7TN89 | GTPase ERas | Reviewed | 230 | MALPTKSSI ... | 3.6.5.2 |
| A0A3G9JYH7 | Luciferase | Reviewed | 271 | MRINISLSS ... | 1.-.-.- |
| ⋮ | ⋮ | ⋮ | ⋮ | ⋮ | ⋮ |

### Technical Validation

In this section, we explore the differences of three datasets, sourcing from DeepFRI^6^, CAFA3^13^, and PAD (reviewed), across different levels of sequence identity. The primary source of data for PAD (reviewed) was the UniProt database^16^, and the curation process involved systematic screening of proteins to select those with comprehensive and expertly reviewed function annotations. We carried out MMseqs2^18^ with default parameters to cluster the sequences of different categories from each dataset at sequence identity thresholds of 10%, 30%, 50%, 70%, and 90%. This is to provide insights into the characteristics of the three datasets.

In evaluating the quality of datasets based solely on the number of sequence clusters, PAD consistently emerges as the superior choice across all the categories (Figure 3). Across various levels of sequence identity, the number of clustered sequences exhibits similar trends. At lower sequence identity thresholds such as 10% and 30%, PAD consistently has better diversity than those of DeepFRI and CAFA3 in terms of the number of sequence clusters. On enzyme commission numbers, PAD consistently demonstrates higher counts of sequence clusters compared to DeepFRI across different sequence identity thresholds (Figure 3A). At lower sequence identities (10% and 30%), PAD consistently outperforms DeepFRI, indicating it has more diverse proteins of enzyme functions. As sequence identity increases, both methods demonstrate a rise in the number of sequence clusters, with PAD exhibiting superior performance compared to DeepFRI. This suggests that PAD is able to provide more diverse proteins across a wide range of sequence identities when training ML methods on predictions of enzyme functions. Across GO terms, PAD consistently outperforms both DeepFRI and CAFA3 in terms of the number of sequence clusters, irrespective of the functional annotations of GO terms (Figure 3B-D). Specifically, Figure 3(B) compares the sequence clusters of biological process (BP) annotated proteins among PAD, DeepFRI, and CAFA3. Similar to the ECN dataset, PAD exhibits superior performance over DeepFRI across all sequence identity thresholds. Notably, at higher sequence identities (70% and 90%), where CAFA3 exceeds DeepFRI in the number of sequence clusters. In Figure 3(C), we observe a consistent trend across the GO BP dataset, where PAD consistently outperforms DeepFRI and CAFA3 across all sequence identity thresholds. Figure 3(D) illustrates this trend by comparing the number of sequence clusters for the MF term. PAD outperforms DeepFRI and CAFA3, indicating its ability to capture a broader diversity of sequences across various sequence identity levels. These findings suggest that PAD offers a more comprehensive representation of sequence space, encompassing a wider range of sequences and functional annotations compared to the other two datasets.

**Figure 3.**
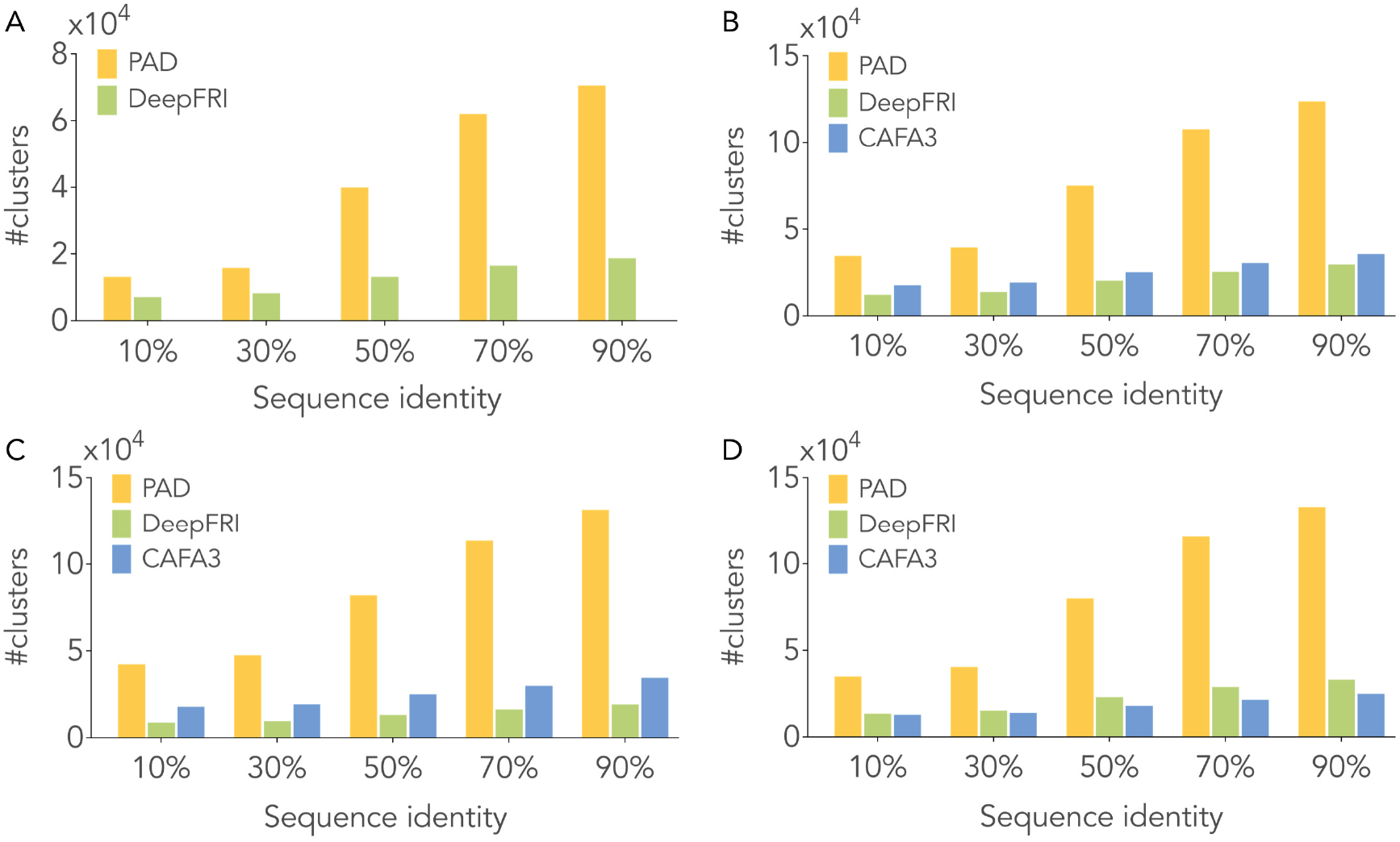
Comparing datasets from PAD, DeepFRI, and CAFA3. Variances in the number of sequence representatives clustered at diverse levels of sequence identity across (A) ECN, (B) BP, (C) CC, and (D) MF categories.

Nevertheless, it is important to acknowledge that the choice of dataset can vary depending on specific research objectives and criteria. While PAD demonstrates strength in quantity by generating a larger number of sequence clusters, DeepFRI and CAFA3 offer distinct advantages in terms of specificity and focus. DeepFRI, for instance, may be preferred in research scenarios where a more refined dataset with fewer clusters is desired, potentially offering higher specificity in terms of sequence identity and functional annotations. Conversely, CAFA3, despite yielding fewer sequence clusters overall, may be more suitable for studies requiring a focused and less redundant dataset, which could facilitate more streamlined analyses and interpretations.

In summary, while PAD excels in collecting a large number of sequences spanning diverse sequence identities and functional annotations, the choice between PAD, DeepFRI, and CAFA3 hinges on specific research objectives and criteria. The comparative analysis reveals clear distinctions in the comprehensiveness and coverage of PAD, DeepFRI, and CAFA3 across various sequence identity thresholds. While PAD emerges as the preferred choice due to its extensive coverage, DeepFRI and CAFA3 provide a reliable alternative for specificity, albeit necessitating supplementation with additional datasets for comprehensive coverage. Researchers should assess their specific needs and the scope of each dataset when selecting the most suitable resource for their development of ML methods for protein function prediction.

### Usage Notes

With the expanding proteins in the UniProt database, machine learning professionals often encounter a scarcity of curated data for predicting protein function, which can hinder their innovation in developing machine learning methods for such tasks. However, precise and prompt screening of proteins with annotated functions is pivotal for developing machine learning methods and validating their ability to generalize to newly sequenced proteins. The PAD dataset is tailor-made to streamline the application of machine learning in protein function annotation tasks. This dataset serves as a valuable resource for various purposes, including: (1) a protein-level multiple-label classification to assign function annotations to proteins with either ECNs or GO terms; and (2) a protein-level binary classification to distinguish enzymes from non-enzymes.

PAD has been curated through an extensive screening process of the UniProt database and is publicly accessible. PAD not only facilitates the training of ML-based algorithms but also provides substantial value for the evaluation and validation of state-of-the-art ML methods. Additionally, it functions as a benchmark for comparing various ML methods dedicated to predicting protein functions. With the release of PAD, our primary goal is to improve the accessibility of ML-based methods for the precise assignment of function annotations to proteins. This initiative not only aims to enhance the predictive performance of these models but also advocates for the open sharing of databases within the field of proteomics. Beyond its direct relevance to our research, this comprehensive dataset makes a contribution to the broader scientific community’s efforts to advance the field of protein function annotation.

There are two main challenges when assigning function annotations to proteins that are not addressed in this dataset. First, unreviewed proteins; we acknowledge that these proteins may have incorrect or partially incorrect labels, potentially introducing bias when training ML models. Although we anticipate that expertly reviewed proteins can help mitigate this bias, it remains speculative and does not account for all the noise stemming from unreviewed proteins. Second, the current dataset does not explicitly address the impact of function annotations filtered by the number of proteins associated with those annotations. As the number of annotations for BP and MF terms is too extensive for ML methods, this presents an important consideration regarding computing memory. For the MF term, there are over 17,000 annotations, posing a significant challenge for ML methods. In summary, the creation of effective datasets represents the initial step in advancing machine learning for the annotation of protein function. Collaborative databases are the preferred approach for developing cutting-edge methods, fostering a genuinely diverse initiative aimed at transforming healthcare

## Footnotes

† [The dataset website](https://kornmann.bioch.ox.ac.uk/jang/services/pad/download.html).

## Notes

### Competing Interest Statement

The authors have declared no competing interest.

## References

1. Abramson, J. et al. Accurate structure prediction of biomolecular interactions with AlphaFold 3. Nature 1–3 (2024).

2. Lin, Z. et al. Evolutionary-scale prediction of atomic-level protein structure with a language model. Science 379, 1123–1130 (2023).

3. Jumper, J. et al. Highly accurate protein structure prediction with AlphaFold. Nature 596, 583–589 (2021).

4. Baek, M. et al. Accurate prediction of protein structures and interactions using a three-track neural network. Science 373, 871–876 (2021).

5. Radivojac, P. et al. A large-scale evaluation of computational protein function prediction. Nat. methods 10, 221–227 (2013).

6. Gligorijevic, V. et al. Structure-based protein function prediction using graph convolutional networks. Nat. communications 12, 3168 (2021).

7. Altschul, S. F. et al. Gapped BLAST and PSI-BLAST: a new generation of protein database search programs. Nucleic acids research 25, 3389–3402 (1997).

8. Pellegrini, M., Marcotte, E. M., Thompson, M. J., Eisenberg, D. & Yeates, T. O. Assigning protein functions by comparative genome analysis: protein phylogenetic profiles. Proc. Natl. Acad. Sci. 96, 4285–4288 (1999).

9. Uetz, P. et al. A comprehensive analysis of protein–protein interactions in Saccharomyces cerevisiae. Nature 403, 623–627 (2000).

10. Yu, T. et al. Enzyme function prediction using contrastive learning. Science 379, 1358–1363 (2023).

11. Ashburner, M. et al. Gene ontology: tool for the unification of biology. Nat. genetics 25, 25–29 (2000).

12. Webb, E. C. et al. Enzyme nomenclature 1992. Recommendations of the Nomenclature Committee of the International Union of Biochemistry and Molecular Biology on the Nomenclature and Classification of Enzymes. Ed. 6 (Academic Press, 1992).

13. Zhou, N. et al. The cafa challenge reports improved protein function prediction and new functional annotations for hundreds of genes through experimental screens. Genome biology 20, 1–23 (2019).

14. Kim, G. B. et al. Functional annotation of enzyme-encoding genes using deep learning with transformer layers. Nat. Commun. 14, 7370 (2023).

15. Jang, Y. J. et al. Accurate prediction of protein function using statistics-informed graph networks. Nat. Commun. (2024).

16. Bateman, A. et al. UniProt: the universal protein knowledgebase in 2023. Nucleic Acids Res. (2022).

17. Stryer L. T. J., Berg JM. Biochemistry (San Francisco: W.H. Freeman, 2002), 5 edn.

18. Steinegger, M. & Söding, J. MMseqs2 enables sensitive protein sequence searching for the analysis of massive data sets. Nat. biotechnology 35, 1026–1028 (2017).

